# Reliable genetic correlation estimation via multiple sample splitting and smoothing

**DOI:** 10.1101/2023.01.15.524097

**Authors:** The Tien Mai

## Abstract

In this paper, we aim to investigate the problem of estimating the genetic correlation between two traits. Instead of making assumptions about the distribution of effect sizes of the genetic factors, we propose the use of a high-dimensional linear model to relate a trait to genetic factors. To estimate the genetic correlation, we develop a generic strategy that combines the use of sparse penalization methods and multiple sample splitting approaches. The final estimate is determined by taking the median of the calculations, resulting in a smoothed and reliable estimate. Through simulations, we demonstrate that our proposed approach is reliable and accurate in comparison to naive plug-in methods. To further illustrate the advantages of our method, we apply it to a real-world example of a bacterial GWAS dataset, specifically to estimate the genetic correlation between antibiotic resistant traits in *Streptococus pneumoniae*. This application not only validates the effectiveness of our method but also highlights its potential in real-world applications.

## Introduction

Genome-wide association studies (GWAS) have been instrumental in demonstrating that various complex traits may be influenced by common genetic variants, as exemplified in studies such as [Giambartolomei et al., 2014, Pickrell et al., 2016, Mancuso et al., 2017]. Recently, there has been a growing interest in understanding the relationship or interaction of common genetic variants across pairs of traits, as seen in studies like [Bulik-Sullivan et al., 2015, Shi et al., 2017, Lu et al., 2017]. This understanding can have a wide range of benefits, such as in epidemiological and etiological studies [Davey Smith and Ebrahim, 2003, Davey Smith and Hemani, 2014], as well as in genetic risk prediction [Purcell et al., 2009, Maier et al., 2015]. By quantifying the interaction of genetic variants across pairs of traits, it can provide useful insights in understanding the underlying genetic mechanisms of the complex traits and can be applied in various fields of genetics.

In the field of genetics, the concept of heritability of a trait has been widely studied and is considered a key quantitative measurement [Lynch and Walsh, 1998, Bürger, 2000]. Building upon this concept, the concept of genetic correlation between two phenotypes has been defined in order to capture the correlation between causal loci [Bulik-Sullivan et al., 2015, Shi et al., 2017, Van Rheenen et al., 2019]. Essentially, this metric is used to measure the extent to which genetic variants across the genome contribute to the correlation between two phenotypes. Understanding the genetic correlation between different traits is informative as it can provide insights into the underlying polygenic genetic architecture of complex traits. It can also be used to identify genetic factors that contribute to the correlation between two traits and helps to understand the underlying genetic mechanisms. Additionally, genetic correlation can also be used to identify pleiotropy, which is when a gene has multiple effects on different traits. Furthermore, the estimation of genetic correlation is an important step in many genetic studies, such as genome-wide association studies (GWAS), as it can help to identify genetic loci that are associated with multiple traits, which can be useful in the study of complex diseases. Moreover, genetic correlation can also be used to identify genetic loci that are associated with the same trait but in different populations. This can provide insights into the underlying genetic architecture of the trait and can help to identify genetic loci that are associated with the trait in different populations.

Traditional approaches to estimating genetic correlation estimation are based on exploring a linear mixed-effect model framework in which the effects of genetic variants are assumed to be random (usually assumed to be normally distributed with 0-mean). This is opposite to the fixed effect assumption embedded in the framework of quantitative genetics theory [Falconer, 1960, Lynch and Walsh, 1998, Lee et al., 2018, Gorfine et al., 2017, Janson et al., 2017, Golan and Rosset, 2018]. Some popular approaches for making inference about genetic correlation in linear mixed model are: maximum likelihood estimation [Loh et al., 2015, Lee et al., 2012, 2013], moment method [Golan et al., 2014, Lu et al., 2017] and linkage disequilibrium score regression [Bulik-Sullivan et al., 2015, Speed and Balding, 2019]. A comprehensive comparison and discussion of these approaches in the context of the random-effect model can be found in a very recent review by [Peyrot et al., 2019].

In this paper, we focus on utilizing a high-dimensional linear regression model to understand the relationship between phenotype and genotype in genetic studies, particularly in the context of genome-wide association studies (GWAS). One of the key advantages of this approach is that it does not rely on any assumptions about the distribution of effect sizes. This is a particularly natural model for investigating the entire genome in GWAS, as it has been shown to be more efficient than traditional univariate methods in GWAS. The use of this model has been previously demonstrated in studies such as [Falconer, 1960, Lynch and Walsh, 1998, Wu et al., 2009, Brzyski et al., 2017, Lees et al., 2020] as a means to improve the understanding of the genetic basis of complex traits. Furthermore, the use of high-dimensional linear regression model allows for the analysis of a large number of genetic variations at once, which can help to identify the most important genetic factors influencing the trait of interest. In this way, we aim to make a contribution to the field by highlighting the benefits of using this model over the traditional univariate approach in GWAS.

Built up on recent advances in machine learning approaches for statistical genetics, we propose an aggregation approach based on multiple sample splittings [Dai et al., 2022, Fei et al., 2019, Fei and Li, 2021]. The main ingredient in our approach is the selective inference framework [Tian, 2020, Tian and Taylor, 2018, Fan et al., 2012, Lee et al., 2016, Tibshirani et al., 2016]. More specifically, it is a two-stage strategy by first splitting samples into two parts, performing variable selection via a sparse regularization (such as Lasso) and then applying partial regression with the other part to provide valid inference under the selected model. This is to ensure that the different latent structures possibly residing in the sample are properly taken into account in both the selection and estimation steps. We obtain a numerically stable smoothed estimates by taking the median of the estimates over multiple random splits.

The idea of “splitting and smoothing” different estimates to yield an estimate with improved statistical properties is the central feature of the generic boosting approach widely used in machine learning, such as AdaBoost [Freund and Schapire, 1996]. It is noted that the estimation based on a single split is highly unstable and thus it is difficult to separate true signals from noises. This has been observed as using a single tree in the bagging algorithm [Bühlmann and Yu, 2002]. In order to reduce this variability, we propose to use a multi-sample splitting scheme that the data is randomly split multiple times and repeat the estimation procedure accordingly. The final estimation is obtained via taking the median of the resulting estimates to obtain the smoothed estimate. The multiple sample splitting approach has previously been proposed in the statistics community, such as in [Meinshausen et al., 2009, Fan et al., 2012], and successfully used in GWAS [Renaux et al., 2020, Buzdugan et al., 2016, Mai et al., 2021].

In order to evaluate the effectiveness of our proposed methods, numerical simulations were conducted. These simulations allowed us to assess the performance of our approach under various conditions and provided valuable insights into its capabilities. Additionally, to further illustrate the utility of our framework, we applied our procedure to a specific case study involving bacterial GWAS. Specifically, we used our methods to estimate the genetic correlations of antibiotic resistant phenotypes in bacteria. This application is particularly noteworthy as, while there has been a considerable amount of research on estimating genetic correlation in human GWAS, the topic has received relatively little attention in the context of bacteria. The results of our case study not only demonstrate the applicability of our methods to this understudied area but also provide valuable insights into the genetic factors underlying antibiotic resistance in bacteria. Overall, the numerical simulations and case study serve to both validate our methods and showcase their potential in real-world applications.

The structure of the paper is as follows: In Section 2, we formally introduce our model and provide a clear and precise definition for the concept of genetic correlation, which is the focus of our study. This section serves as a foundation for the rest of the paper and sets the stage for the subsequent sections. In Section 3, we delve deeper into the proposed method and provide a detailed explanation of the steps involved in the analysis. This section is of particular importance as it presents the core of our work and the key contributions of the paper. The methods are explained in a clear and easy-to-follow manner and are supplemented with relevant mathematical derivations, where appropriate. To demonstrate the effectiveness and practical utility of our method, we conduct numerical studies with simulations and a real data application in Section 4. This section provides a thorough evaluation of our method and serves to validate its performance under various scenarios. Finally, in the last section, we provide a discussion of the results, highlighting the key findings of our study and their implications. We also provide a conclusion that summarizes the main contributions of our paper and highlights the potential future directions of research in this area.

## Model

We study the problem of genetic correlation estimation based on individual-level GWAS data. More specifically, we observe two traits *y* and *z* of *n* samples and a genotype matrix *X* of size *n* × *p* (often SNPs). Each trait is modelled as a linear combination of *p* genetic variants *X*_.*j*_ and an error term (environmental and unmeasured genetic effects)

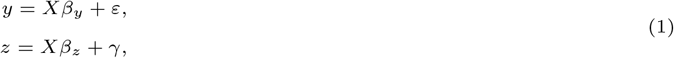

where *β*_*y*_, *β*_*z*_ are vectors of SNP effect sizes of length *p*. We assume that *X*_*i*._ are i.i.d random vector with 0-mean and covariance matrix Σ and that the random noises *ε*_*i*_, *γ*_*i*_ are independent of *X* with 0-mean and the variances 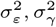.

In this study, a key departure from the typical approach in the field is the lack of assumptions regarding the distribution of effects *β*_*y*_ and *β*_*z*_. Most research in this area, particularly within the framework of linear mixed models, relies on such assumptions to inform the analysis. However, by forgoing these assumptions, our work allows for a more flexible and inclusive approach to understanding the underlying relationships. This unique perspective may provide new insights and better understanding of the data. Additionally, this method can be useful in case of non-normally distributed data and can be less restrictive. This can be especially beneficial in cases where the distribution of the effects may not be known or may be difficult to accurately estimate. Overall, this approach provides a fresh perspective on the data, and can yield new insights into the underlying relationships being studied.

### Genetic correlation

From the problem formulation (1), the covariance between the phenotypes *y* and *z* can be explained as a summation of the genetic covariance and environmental covariance as follows

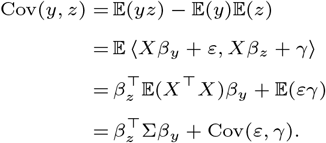

As a consequence, the *genetic correlation* between two traits *y* and *z* is defined by

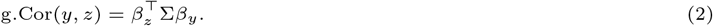

This can then be further normalized to obtain the genetic correlation scaling between −1 and 1; where −1 indicating perfect negative correlation and 1 showing perfect positive correlation. More precisely, we define

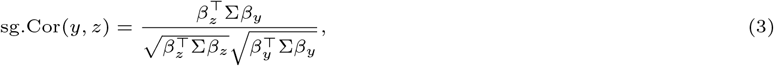

which provides a standardized genetic correlation scale between −1 and 1 and thus, can be used to compare across pairs of traits and it is possible to compare across genomic regions.

### Plug-in Lasso type estimation

We can obtain naive estimates of genetic correlation (2) and its normalization (3) by using some estimates 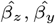 of the effect sizes and 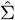 of the covariance matrix. More specifically, by using the Lasso method, one can obtain the non-zero estimated effect sizes of the selected covariates, and one can also use these covariates to obtain a sample covariance matrix. More precisely, let 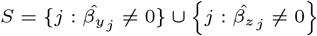 where 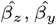 are estimates from the Lasso method,

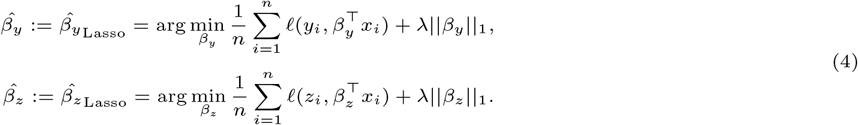

Here *ℓ*(*a, b*) is the negative log-likelihood for an observation e.g. for the linear Gaussian case it is 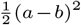 and for logistic regression it is −*a* · *b* +log(1 + *e*^*b*^). The tuning parameter *λ >* 0 controls the overall strength of the penalty and we use 10-fold cross-validation to choose a suitable value for *λ*. The Lasso is implemented in the `R` package `glmnet` [Friedman et al., 2010].

Now, we can calculate the quantity of interest by plugging-in these Lasso estimates, with 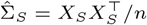,

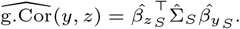

Other quantities can also be estimated directly via plugging-in Lasso outputs.

However, as shown in simulations in Section 4, these naive plug-in approaches would create a huge bias. Moreover, uncertainty quantification for this kind of approach is difficult to obtain and actually not known up to present. To overcome these problems, we next present a reliable approach via multiple sample splitting and aggregation which provides an accurate estimate together with some reliable uncertainty quantifications.

## Method

In this section, we introduce our generic strategy to estimate genetic correlation, which consists of a multi sample splitting strategy and a sparse regularization step followed by an estimation step.

### Estimating via multiple sample splitting and aggregation

First, without loss of generality, let us assume that the sample size is even for simplicity. The original dataset (*y, z, X*) is randomly divided into two disjoint datasets 𝒟_1_ = (*y*^(1)^, *z*^(1)^, *X*^(1)^) and 𝒟_2_ = (*y*^(2)^, *z*^(2)^, *X*^(2)^) with equal sample sizes. The sample splitting is a useful approach that can help to eliminate overfitting when variable selection and subsequent estimation is performed on the same dataset [Fan et al., 2012, Buzdugan et al., 2016, Li et al., 2019].

First, we apply a variable selection step on data 𝒟_1_ as in (4), where we propose using Lasso as a default alternative, to select the most relevant covariates (to reduce dimension of the model). Other variable selection methods could be used, for example, the sure independence screening (SIS) procedure [Fan and Lv, 2008]. Denote *S*_*y*_ ⊂ {1, …, *p*} and *S*_*z*_ ⊂ {1, …, *p*} the subset of important predictors obtained respectively for the trait *y* and *z*, where |*S*_*y*_| *< n/*2 and |*S*_*z*_| *< n/*2.

Then, using data 𝒟_2_, we fit linear regression on 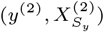 and 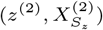 to obtain unbiased estimates of the regression coefficients *β*_*y*_ and *β*_*z*_. The estimation of genetic correlation between two traits is calculated as in (2) where the sample covariance matrix is used. Thus, it is guaranteed that variable selection and estimation are performed on independent samples.

Note that, in order to stay away from depending crucially on a particular sample splitting employed, we propose to perform the sample splitting and inference procedure many times (e.g. 100 times) and aggregating the corresponding results via taking the median. Our proposed strategy is summarized in Algorithm 1. Another important point in our proposed method, that is different to other works, is to aggregate the final result via the median which was inspired from reference [Lugosi and Mendelson, 2019]. This is further confirmed from our simulation studies in Section 4.

#### Algorithm 1 B.CORE algorithm

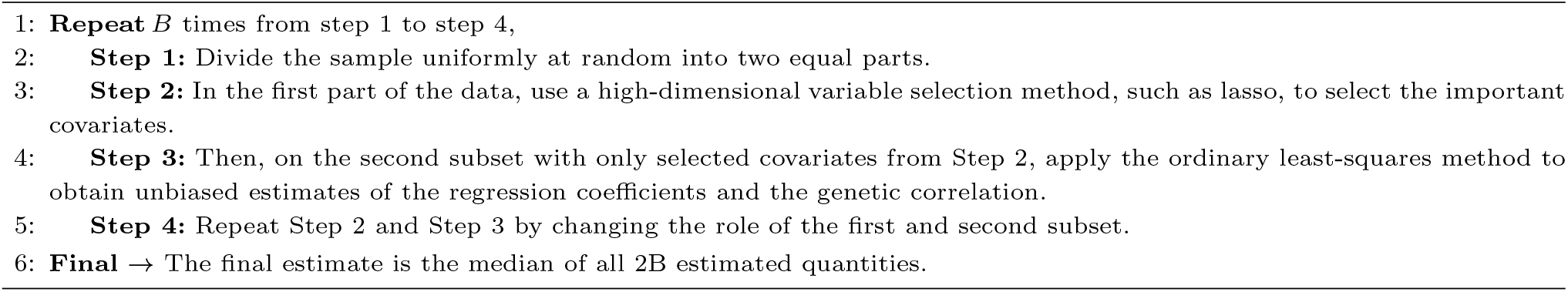

In addition, switching the roles of the data subsets in **Step 4** helps us to obtain another estimation and thus would lead to a more stable estimation. It is noted that the main cost for our Algorithm 1 is in fitting a penalized regression (**Step 2**) for variable selection in the setting where *p* > *n*. It is, however, important to note that there have been recent advancements in methods for quickly analyzing large GWAS data using penalized regression, see e.g [Qian et al., 2020]. Additionally, the process can be easily made more efficient by implementing multiple repetitions in parallel.

Also, as suggested by the work in [Gorfine et al., 2017], we could also reduce the ultra-high dimensionality to a relatively large scale before applying Algorithm 1 by using a screening method [Fan and Lv, 2008] for example: a marginal-type screening technique such as SIS (as long as the number of SNPs involved is not large, e.g., 10,000), for higher numbers of SNPs, a joint-type screener, such as the ITRRS, should be used so that the LD between the SNPs is considered and the truly associated SNPs can be better selected.

Theoretical foundations for using the splitting and smoothing/aggregation method to estimate regression coefficients have been established in previous research for both linear models (as outlined in [Fei et al., 2019]) and generalized linear models (as in [Fei and Li, 2021]).

### A 95% reliable interval

It is important to note that by repeating sample splitting, we are able to obtain a range of estimates for the quantity of interest. This is beneficial because it allows us to construct a meaningful interval of the estimated values. More specifically, by using the outputs from Algorithm 1, which is 2*B* estimated quantities of interest, we can construct a 95% reliable interval by taking the interval from the 2.5% to 97.5% quantiles of the outputs. This is an important feature of our proposed method, as it allows us to not only estimate the genetic correlation, but also to quantify the uncertainty of the estimate. It is well-known that interval estimates are more informative than point estimates, as they provide a range of plausible values for the true parameter.

The simulation results presented in Section 4 provide further evidence of the accuracy of our proposed method. These simulations show that the 95% reliable interval is highly accurate, in the sense that it always contains the true value of interest. This is a strong indication that our proposed method is able to provide accurate and reliable estimates of genetic correlation.

Moreover, by providing interval estimates, our method allows for a more comprehensive understanding of the underlying relationships, as it allows for a more thorough assessment of the uncertainty of the estimates. This is particularly important when dealing with complex traits, where the underlying distribution of effects may not be easily identifiable or may not conform to a known distribution. This feature of our method provides a more robust and inclusive approach to understanding.

### Extension with traits on different samples

We now describe a model in the general setting where the traits are not observed on the same set of samples. More specifically, two traits *y* and *z* are collected from two different sample sets of *n*_1_ and *n*_2_ sizes, each is modelled as a linear combination of *p* genetic variants *X*_.*j*_ and an error term (environmental and unmeasured genetic effects) as following

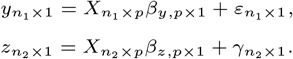

The Algorithm 1 can be used to estimate the genetic correlation between these two traits. More specifically, the multiple sample splitting will be conducted on each sample set to estimate the effect size and the sample covariance matrix could be estimated by merging these sample set.

### Numerical studies

#### Simulation studies

##### Experimental designs

In this work, we use a real data set of 3069 *Streptococus pneumoniae* genomes that was collected from an infant cohort study conducted in a refugee camp on the Thailand-Myanmar border [Chewapreecha et al., 2014, Lees et al., 2016]. This dataset provides us with a unique opportunity to study the genetic correlation of antibiotic resistant traits in bacteria and to create semi-synthetic datasets that incorporate the levels of population structure and linkage disequilibrium that are present in natural populations. As seen in Figure 1, we use a fully observed genotype matrix of 3051 samples and 5000 SNPs to conduct our analysis, which allows us to gain a better understanding of the genetic factors that are associated with antibiotic resistance in *Streptococcus pneumoniae*.

**Fig. 1.**
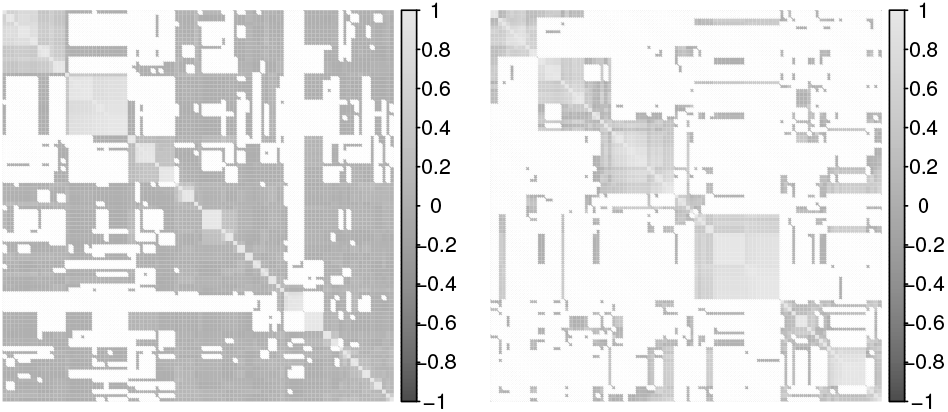
Sample correlation matrices of the 100 random SNPs (right) and 100 samples (left) in the genotype matrix displays the complex dependence structure presented in the real Maela data.

Given the genotypes, we consider the following settings for choosing the causal SNPs (non-zero effect sizes) under varying genetic architectures and the phenotypes are simulated as in model (1) :

Setting I: genetic basis overlap, i.e. *β*_*z*_ and *β*_*y*_ have 50% non-zeros components in common. Then we have the following scenario:

- (Ia). *y* and *z* are polygenic that *β*_*z*_ and *β*_*y*_ are with 1000 non-zero components.
- (Ib). *y* and *z* are sparse that *β*_*z*_ and *β*_*y*_ are with 50 non-zero components.
- (Ic). *y* is polygenic that *β*_*z*_ is with 1000 non-zero components, while *z* is sparse that *β*_*y*_ is with 50 non-zero components.

Setting II: genetic basis non-overlap, i.e. *β*_*z*_ and *β*_*y*_ have no common non-zeros components. Then we have the following scenario:

- (IIa). *y* and *z* are polygenic that *β*_*z*_ and *β*_*y*_ are with 1000 non-zero components.
- (IIb). *y* and *z* are sparse that *β*_*z*_ and *β*_*y*_ are with 50 non-zero components.
- (IIc). *y* is polygenic that *β*_*z*_ is with 1000 non-zero components, while *z* is sparse that *β*_*y*_ is with 50 non-zero components.

Given the above setting for choosing causal SNPs, the non-zero coefficients of *β*_*z*_ and *β*_*y*_ are drawn from the normal distribution 𝒩 (0, 1). The noise follows the normal distribution 𝒩 (0, 1). We further vary the size of the samples by uniformly subsampling the samples between 1000, 2000 or full samples. For each setup, we generate 50 simulation runs and report the mean and the standard deviation of the absolute errors for each method across the simulation runs.

We performed simulations to compare the accuracy of our proposed method to other estimators of genetic correlation under different genetic architectures. We compare Lasso, B.CORE (mean) and B.CORE (median). The number of repeated sample splitting is performed with *B* = 50 times. The Lasso is used with 10-fold cross validation for choosing the tuning parameter *λ*.

##### Simulation results

As demonstrated in the simulation results presented in Figure 5, 6, 7 and 8, our proposed method of aggregating the outputs from multiple sample splitting using the median is more accurate than using the mean. This finding is expected, as our target here is a correlation quantity [Lugosi and Mendelson, 2019] and, also, for the real dataset, the samples would no longer be independent. Additionally, we also found that the median-aggregation leads to a much smaller standard deviation compared to the mean. This is an important aspect, as a smaller standard deviation indicates that the results are more consistent and less variable, making the estimates more reliable.

This conclusion is also supported by the fact that when we look at the distribution of the estimates obtained from both the median and mean aggregation, the median aggregation yields more precise and concentrated estimates around the true value, while the mean aggregation yields estimates that are more dispersed. Moreover, we are not assuming any specific underlying distribution for the data and hence the median is a more robust statistic. It is less sensitive to outliers and skewness in the data, which can be present in real datasets. In addition, it is worth noting that our proposed method allows for a more flexible and inclusive approach to understanding the underlying relationships. This is particularly important when dealing with complex traits, where the underlying distribution of effects may not be easily identifiable or may not conform to a known distribution.

In comparison with the plug-in Lasso method, we found that our approach quite often yields better results. This is more often to be the case where the same size is small. Most importantly, our approach returns interval estimations which allow to obtain, for example, confident intervals while it is challenging to obtain such results for the Lasso method. The standard deviation for Lasso and the mean-aggregation seem comparable.

In order to determine the optimal number of sample splittings, *B*, to use in practice, we conducted a series of simulations where we varied *B* across different simulation settings. The results, presented in Figure 2, show that the number of sample splitting can greatly affect the estimation results in various settings. As seen from the figure, the accuracy of the estimates improves as the number of sample splittings increases, but it reaches a plateau after a certain point. This is because, as the number of sample splittings increases, the estimation becomes more precise and the variability of the estimates decreases, but at some point, additional sample splittings would not provide much improvement.

**Fig. 2.**
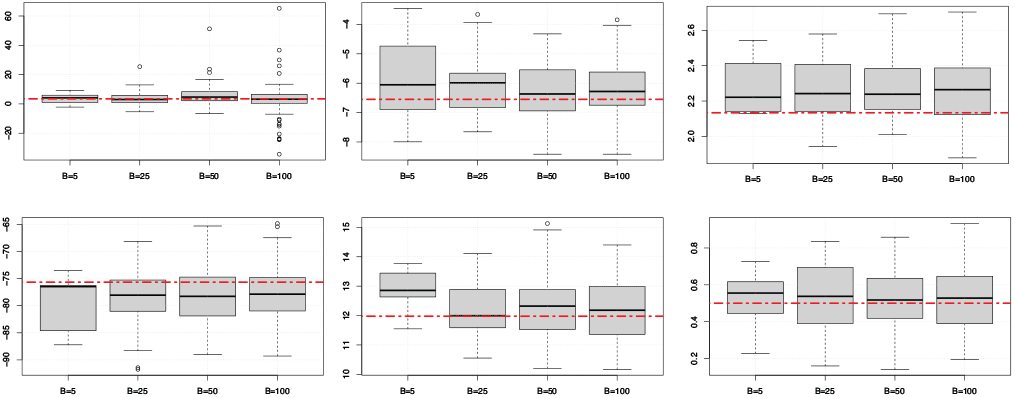
Simulation results for varying the number of sample splitting *B* (5,25,50,100) in different settings. Top row plots are for Setting I; bottom row plots are for Setting II.

Based on the results of our simulations, a value of *B* = 50 is recommended for practical use. This value strikes a balance between computation time and accuracy. However, when computational resources permit, a larger number of *B* can be used to achieve even more stable results. For example, a larger number of *B* will lead to more precise estimates, which can be useful in situations where the sample size is small or the correlation is weak. Additionally, a larger number of *B* can also be useful when there is a high degree of uncertainty in the data, as it allows for a more robust estimation of the genetic correlation.

In addition to evaluating the effectiveness of our proposed method, we also conducted experiments to investigate the effect of different sample splitting sizes. In particular, we varied the proportion of samples allocated to each split (*α*) and evaluated the performance of our method under different conditions. The results, presented in Figure 3, show that the minimum mean absolute error was achieved when *α* = 0.5, suggesting that equal-size splitting is a rational choice in practice. This outcome can be explained by the fact that when the sample size is not large enough, the estimation of genetic correlation is sensitive to the size of sample splitting, and equal-size splitting is a practical choice. This is because equal-size splitting ensures that each split has enough sample size and thus the estimation is more stable and accurate. Furthermore, equal-size splitting also ensures that the sample splitting is random and unbiased, which is crucial for the validity of our inference. Additionally, it is worth noting that equal-size splitting is also computationally efficient. As the sample size increases, the computation time also increases, and equal-size splitting can balance the computation time. This is important when dealing with large datasets and computational resources are limited.

**Fig. 3.**
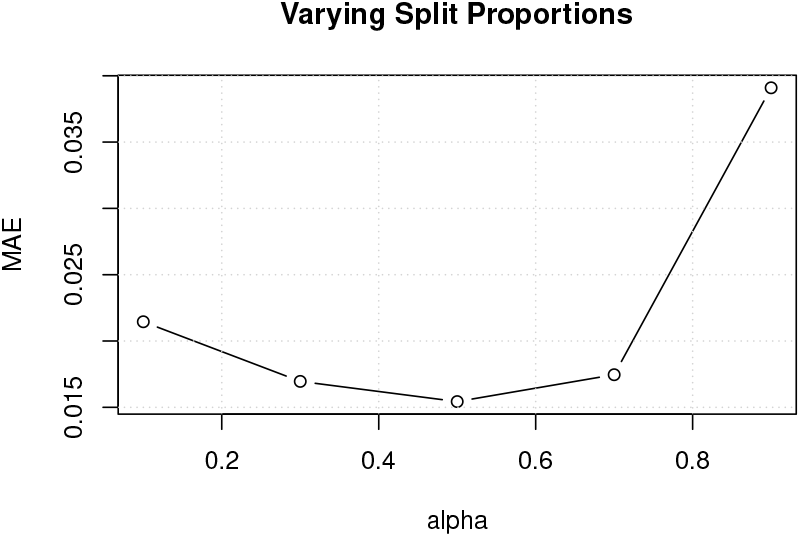
Average mean absolute error (MAE) of the median predictor at split proportions alpha’s (*α*) from 0.1 to 0.9.

## Real data application

In this study, we aimed to examine the genetic correlations between resistance to different antibiotics in *Streptococcus pneumoniae*. To accomplish this, we used the Maela data set, which is a collection of 3069 Streptococcus pneumoniae genomes from an infant cohort study conducted in a refugee camp on the Thailand-Myanmar border. This dataset has been previously used in genome-wide association studies to identify genetic loci associated with antibiotic resistance [Chewapreecha et al., 2014, Lees et al., 2016]. We began by applying standard population genomic procedures to the data, such as using a minor allele frequency threshold and removing missing data, to obtain a genotype matrix with 121014 SNPs. We then considered resistances to five different antibiotics (Chloramphenicol, Erythromycin, Tetracycline, Penicillin and Co-trimoxazole) as the phenotypes.

The results of our analysis are presented in Figure 4. Our findings indicate that there were small to moderate genetic correlations between Chloramphenicol resistance and each of the other antibiotics considered. However, we observed higher genetic correlations between Erythromycin resistance, Tetracycline resistance and Co-trimoxazole resistance. Notably, the most significant genetic correlation was found to be between Penicillin resistance and Co-trimoxazole resistance, with a correlation coefficient of 0.45 (95% CI: 0.40-0.49).

**Fig. 4.**
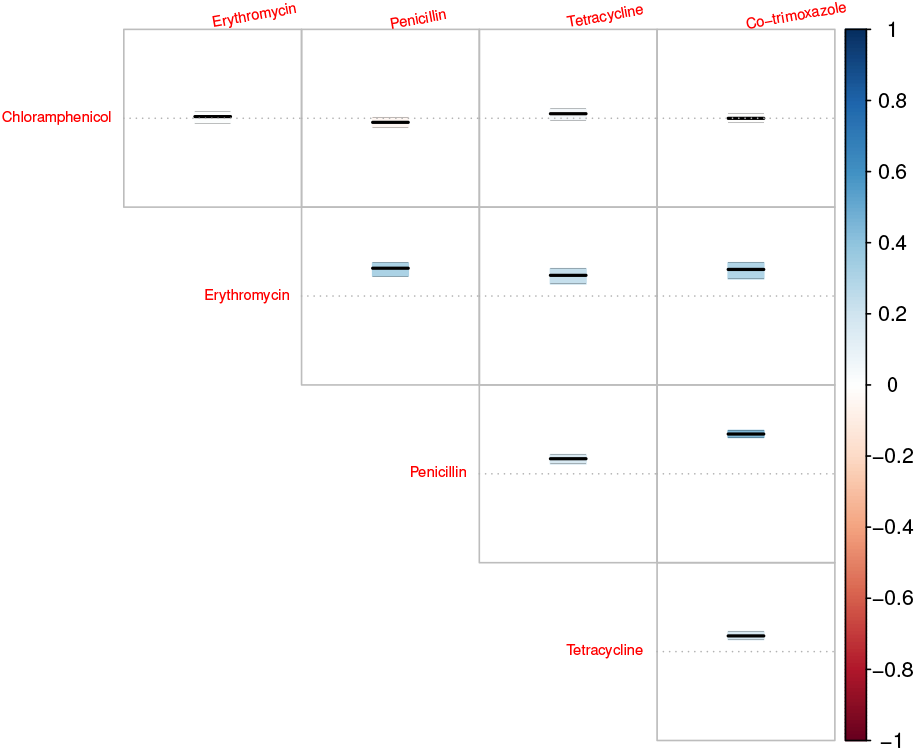
Estimation results of the normalized genetic correlation between five different antibiotic resistances in Maela real dataset.

**Fig. 5.**
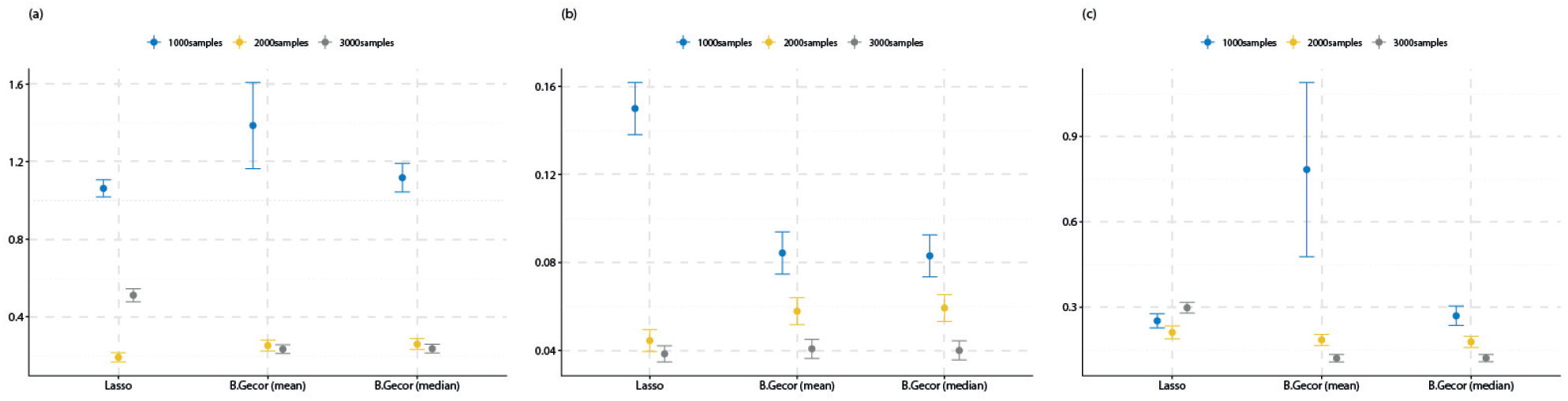
Simulation results on the absolute errors in estimating genetic correlation for Setting I with overlapping genetic basis between the traits.

**Fig. 6.**
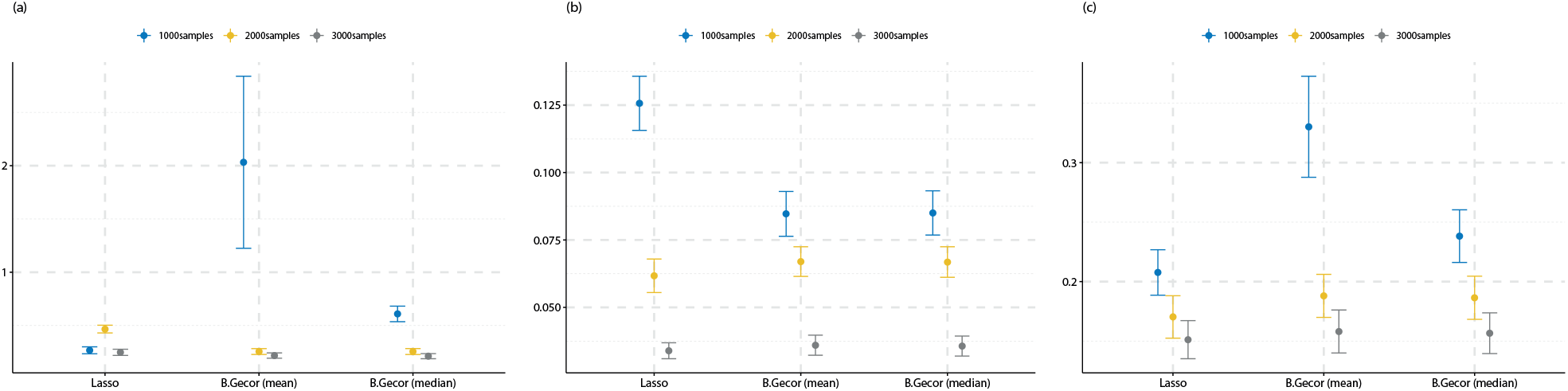
Simulation results on the absolute errors in estimating genetic correlation for Setting II with non-overlapping genetic basis between the traits.

**Fig. 7.**
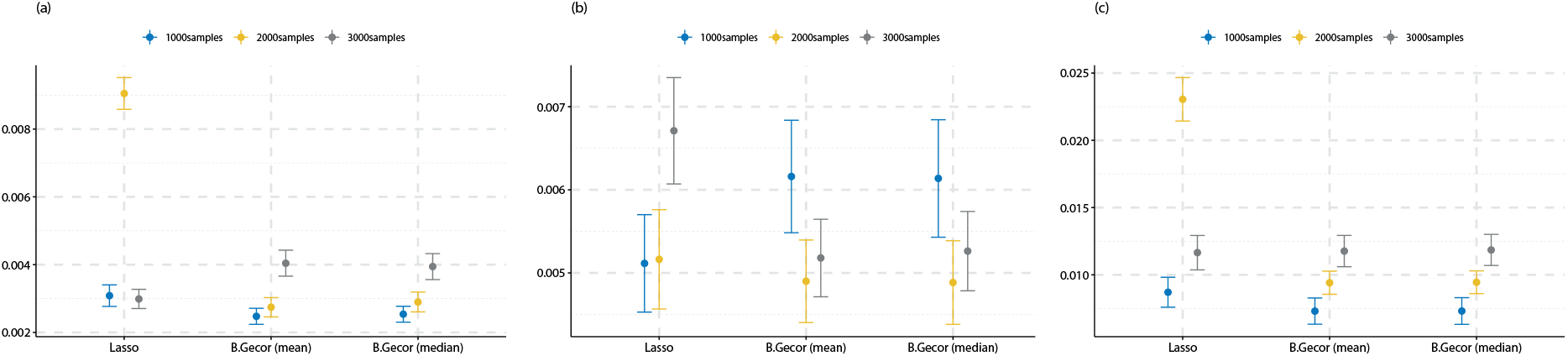
Simulation results on the absolute errors in estimating the standardized genetic correlation for Setting I with overlapping genetic basis between the traits.

**Fig. 8.**
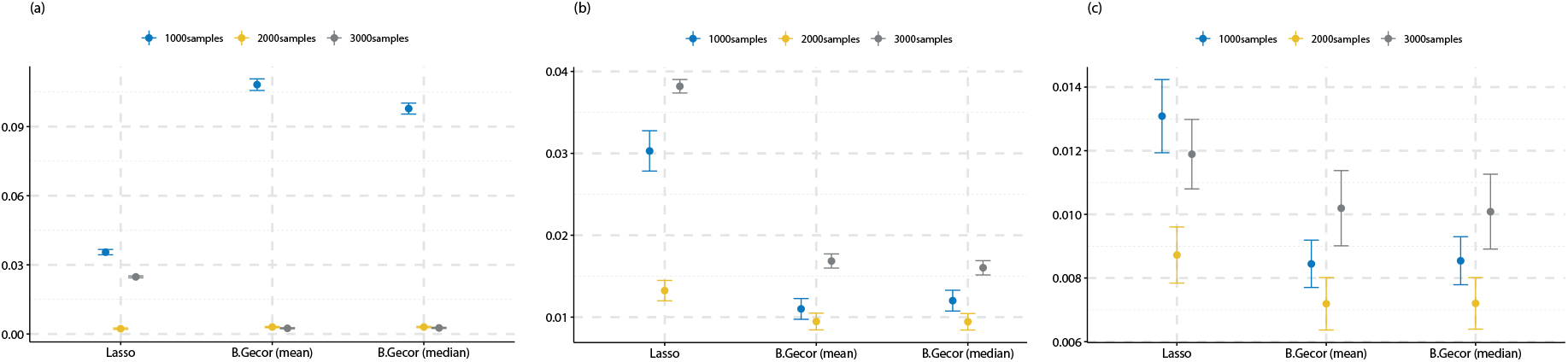
Simulation results on the absolute errors in estimating the standardized genetic correlation for Setting II with non-overlapping genetic basis between the traits.

Our results provide valuable insights into the genetic factors underlying antibiotic resistance in Streptococcus pneumoniae. They also demonstrate the potential of our proposed approach in identifying genetic correlations between different traits in real-world datasets. Furthermore, these results can be used to understand the underlying genetic mechanisms of antibiotic resistance in Streptococcus pneumoniae and could be useful in the development of new treatments and strategies to combat antibiotic resistance.

## Conclusion

In conclusion, genetic correlation is a vital concept in the field of genetics that allows us to understand the relationship between different traits and identify genetic loci that are associated with multiple traits. It is a powerful metric that provides valuable insights into the underlying genetic architecture of complex traits and has been widely studied under various models to better understand the correlation between causal loci.

Estimating genetic correlation is a challenging statistical problem that is gaining increasing attention from researchers in various fields. In this work, we have presented a novel computational strategy for estimating genetic correlation efficiently and accurately. Our approach does not rely on any assumptions about the distribution of effect sizes, allowing for a more flexible and inclusive approach to understanding the underlying relationships. Furthermore, we have demonstrated the efficacy of our method through numerical simulations and an application to a real dataset, where we estimated the genetic correlation between antimicrobial resistant traits in Streptococcus pneumoniae.

Overall, our proposed method is a valuable tool for researchers studying genetic correlation and can be applied to a wide range of complex traits and populations. It is an important step forward in understanding the underlying genetic mechanisms of complex traits and can provide valuable insights into the genetic basis of different diseases.

## Acknowledgments

TTM is supported by the Norwegian Research Council grant number 309960 through the Centre for Geophysical Forecasting at NTNU.

## Availability of data and materials

The R codes and data used in the numerical experiments are available at: https://github.com/tienmt/bcore.

